# Tandemly duplicated *MYB* genes specifically in the Phaseoleae lineage are functionally diverged in the regulation of anthocyanin biosynthesis

**DOI:** 10.1101/2023.07.15.549139

**Authors:** Ruirui Ma, Wenxuan Huang, Quan Hu, Guo Tian, Jie An, Ting Fang, Jia Liu, Jingjing Hou, Meixia Zhao, Lianjun Sun

## Abstract

Gene duplications have long been recognized as a driving force in the evolution of genes, giving rise to novel functions. The soybean genome is characterized by a large extent of duplicated genes. However, the extent and mechanisms of functional divergence among these duplicated genes in soybean remain poorly understood. In this study, we revealed that tandem duplication of *MYB* genes, which occurred specifically in the Phaseoleae lineage, exhibited a stronger purifying selection in soybean compared to common bean. To gain insights into the diverse functions of these *MYB* genes in anthocyanin biosynthesis, we examined the expression, transcriptional activity, metabolite, and evolutionary history of four *MYB* genes (*GmMYBA5*, *GmMYBA2, GmMYBA1* and *Glyma.09g235000*), which were presumably generated by tandem duplication in soybean. Our data revealed that *Glyma.09g235000* had become a pseudogene, while the remaining three *MYB* genes exhibited strong transcriptional activation activity and promoted anthocyanin biosynthesis in different soybean tissues. Furthermore, *GmMYBA5* produced distinct compounds in *Nicotiana benthamiana* leaves compared to *GmMYBA2* and *GmMYBA1* due to variations in their DNA binding domains. The lower expression of anthocyanin related genes in *GmMYBA5* resulted in lower levels of anthocyanins compared to *GmMYBA2* and *GmMYBA1*. Metabolomics analysis further demonstrated the diverse and differential downstream metabolites, suggesting their functional divergence in metabolites following gene duplication. Together, our data provided evidence of functional divergence within the *MYB* gene cluster following tandem duplication, which shed light on the potential evolutionary direction of gene duplications during legume evolution.

## Introduction

In flowering plants, polyploidy or whole-genome duplication is a key process that leads to gene duplication, which provide genetic resources for generating functional novelty (Soltis et al., 2015; Soltis and Soltis, 2016). In addition to polyploidy, another two important processes that result in an increase in gene copy number are segmental and tandem duplication (Rizzon et al., 2006; Freeling, 2009). Following duplication events, duplicated genes undergo functional divergence, which can occur through pseudogenization resulting in loss of function, neofunctionalization leading to the acquisition of novel functions, or subfunctionalization where the duplicated genes retain partial functions of the ancestor gene (Sandve et al., 2018). The phenomenon of functional divergence in duplicated genes has been observed in many species. For instance, in citrus, the *Ruby2*–*Ruby1* gene cluster exhibits subfunctionalization, with these two genes exerting opposite effects in the regulation of anthocyanin biosynthesis (Huang et al., 2018). In maize, the duplicated *MYB* genes, *P1* and *P2*, show different expression patterns that contribute to tissue-specific pigmentation (Zhang et al., 2000). In *Solanum commersonii*, the tandem paralogs *ScAN1* and *ScAN2* have diverged in function with one gene specialized in anthocyanin production and the other one maintaining the conserved function of responding to cold stress (D’Amelia et al., 2018). These examples clearly demonstrate the functional divergence of genes following duplication events. Soybean, which has undergone two polyploidy events, has experienced substantial tandem duplication events, resulting in a significant expansion of genes involved in the anthocyanin biosynthetic pathway (Kim et al., 2012).

Anthocyanins, one of the largest groups of plant flavonoid compounds, not only confer appealing colors to plants but also contribute their tolerance to biotic and abiotic stresses, including drought, cold, ultraviolet (UV)-B, heavy metals, herbivores, and pathogens (Gould, 2004; Hichri et al., 2011; Kovinich et al., 2015). Moreover, anthocyanins offer human health benefits to by protecting against chronic diseases such as metabolic syndrome, cardiovascular disease, and certain cancers (Zhang *et al*., 2014; Putta et al., 2017). The biosynthesis of anthocyanins stems from the general phenylpropanoid pathway and involves several catalytic enzymes, including chalcone synthase (CHS), chalcone isomerase (CHI), flavanone 3-hydroxylase (F3H), flavonoid 3’hydroxylase (F3’H), flavonoid 3’5’ hydroxylase (F3’5’H), dihydroflavonol 4-reductase (DFR), anthocyanidin synthase (ANS), and UDP-flavonoid glucosyltransferase (UFGT) (Sundaramoorthy et al., 2015; Xu et al., 2015) (Supplementary Fig. S1). The expression of anthocyanin biosynthetic genes is primarily regulated by the MYB-bHLH (helix-loop-helix)-WD repeat (MBW) transcriptional complex (Albert et al., 2014; Lloyd et al., 2017; Wang et al., 2021; Wang et al., 2022). The critical part of the MBW transcriptional complex is MYB transcription factors, which are the key determinants of pigmentation and have been identified in various crops such as maize, rice, wheat and foxtail millet (Grotewold et al., 1994; Chin et al., 2016; Shin et al., 2016; Li et al., 2022b). Specific patterns and spatial localizations of anthocyanins determined by MYB transcription factors have also been reported in apple, snapdragon, petunia and lily (Liu et al., 2015). However, limited research has focused on the diverse metabolites and anthocyanin components, which play distinct roles in environmental adaptation produced by tandemly duplicated *MYB* genes.

Soybean (*Glycine max* (L.) Merr.) is one of the world’s most important legume crops that provides plant protein, oil and other essential ingredients for humans and livestock (Malle *et al*., 2020). Compared to its wild relative *Glycine soja* (Sieb. & Zucc.), cultivated soybean and landraces exhibit a wide range of morphological types in order to meet human needs (Li et al., 2008). Various parts of the soybean plant, including hypocotyls, petioles, flowers, and seeds display significant natural variation in color due to distinct accumulation and distribution patterns of anthocyanins and proanthocyanins (Jeong et al., 2019; Xie et al., 2019). Previous studies have shown that the loss of anthocyanin pigment in cereals during domestication, diversification, and improvement is controlled by the same *MYB* gene (Li et al., 2022b). However, only a limited number of transcription factors responsible for anthocyanins biosynthesis have been studied in legume (Lu et al., 2021). The function and evolutionary history of syntenic block in legume is unknown. Therefore, it is crucial to investigate the mechanisms underlying the molecular mechanisms and functional divergence of tandemly duplicated MYB transcription factors in the regulation of anthocyanin biosynthesis in legumes.

Previous studies have described the collinear region of the MYB transcription factors cluster in legumes, which conservatively contributed to seed coat color in soybean, common bean and cowpea (Zabala et al., 2014; García-Fernández et al., 2021; Herniter et al., 2018). In this study, we revealed the tandemly duplicated *MYB* gene cluster that emerged prior to the divergence of soybean from other legume species during the common legume teraploidy event. We observed a stronger purifying selection acting on this cluster in soybean. With the exception of one pseudogene (*Glyma.09g235000*), all duplicated genes exhibited the potential to activate anthocyanin biosynthesis with tissue-specific patterns. Through ectopic expression and metabolomics analysis, we found that the *MYB* genes exhibited functional divergence in activation activities and specificity of downstream target catalytic enzymes, and metabolites. Additionally, a domain swap experiment suggested the divergent R2R3 DNA binding domain primary accounted for the distinct phenotype observed in *GmMYBA5*. Metabolomics analysis further demonstrated the diverse and differential downstream metabolites, suggesting their functional divergence in metabolites following gene duplication. Taken together, our findings shed light on the characteristics of functional divergence within a *MYB* gene cluster, and provide new insights into the evolutionary history of gene duplications and functional divergence as a mechanism driving adaptation during legume evolution.

## Results

### Tandem duplication of the *MYB* genes occurs specifically in the Phaseoleae lineage and exhibits a stronger purifying selection in soybean

Soybean has undergone two polyploidy events and substantial tandem duplication events, resulting in a significant expansion of duplicated regions and genes involved in the regulation of anthocyanin biosynthesis, specifically the MYB transcription factors (Kim et al., 2012). The *R* locus (*GmMYBA2*, *Glyma.09g235100*), is an R2R3 MYB activator located on chromosome 9. It is flanked by three tandemly duplicated *MYB* genes: *GmMYBA5* (*Glyma.09g234900*), *GmMYBA1* (*Glyma.09g235300*) and *Glyma.09g235000* (Gao et al., 2021) (Fig. 1A). *Glyma.09g235000* was identified as a pseudogene due to its lack of expression in any soybean tissue (Gillman et al., 2011). To shed light on the evolutionary forces driving the divergence of the *MYB* genes, we analyzed a region containing the *MYB* gene cluster in different legumes genomes, which have been known to contributed conservatively to seed coat color in soybean, common bean, and cowpea (Zabala et al., 2014; García-Fernández et al., 2021; Herniter et al., 2018). The results indicated that the numbers of tandemly duplicated *MYB* genes varied across different species (Fig. 1B). In certain species such as *Trifolium pratense*, *Cicer arietinum*, *Pisum sativum* and *Aeschynomene evenia*, which diverged earlier from the legume common tetraploidy event, there was only a single copy of the *MYB* transcription factor gene in the collinear regions (Wang et al., 2017). However, an increased number of *MYB* genes was observed in the genera of *phaseolus* and *vigna*, which diverged from soybean approximately 27.3 million years ago (MYA). Notably, we detected four homologous genes in the syntenic region of common bean (*Phaseolus vulgaris*) (Fig. 1B). Interestingly, although all four homologous genes were present, the best matches for *GmMYBA5*, *GmMYBA2* and *GmMYBA1* in common bean were found to be *Phvul.008G038200*. To further explore the evolutionary divergence of these three soybean MYB genes, we next compared the evolutionary distance of the three soybean *MYB* genes using common bean as an outgroup. Our data revealed that ω (Ka/Ks, where Ka represents nonsynonymous substitution and Ks represents synonymous substitution) for *GmMYBA5* was higher than that of *GmMYBA2* and *GmMYBA1* (Table 1), suggesting that *GmMYBA5* has experienced a lower intensity of purifying selection compared to the other two genes. Furthermore, our data indicated that the *MYB* genes in soybean exhibited lower rates of substitution for both Ka and Ks when compared to common bean, consistent with the genome-wide analysis of all genes between soybean and common bean (Zhao et al., 2017). Intriguingly, despite the significantly higher ω observed at the genome-wide level in soybean (Zhao et al., 2017), a lower ω was observed specially for these *MYB* genes in soybean, suggesting that the *MYB* genes have undergone a stronger purifying selection in soybean after their split with common bean.

**Figure 1.**
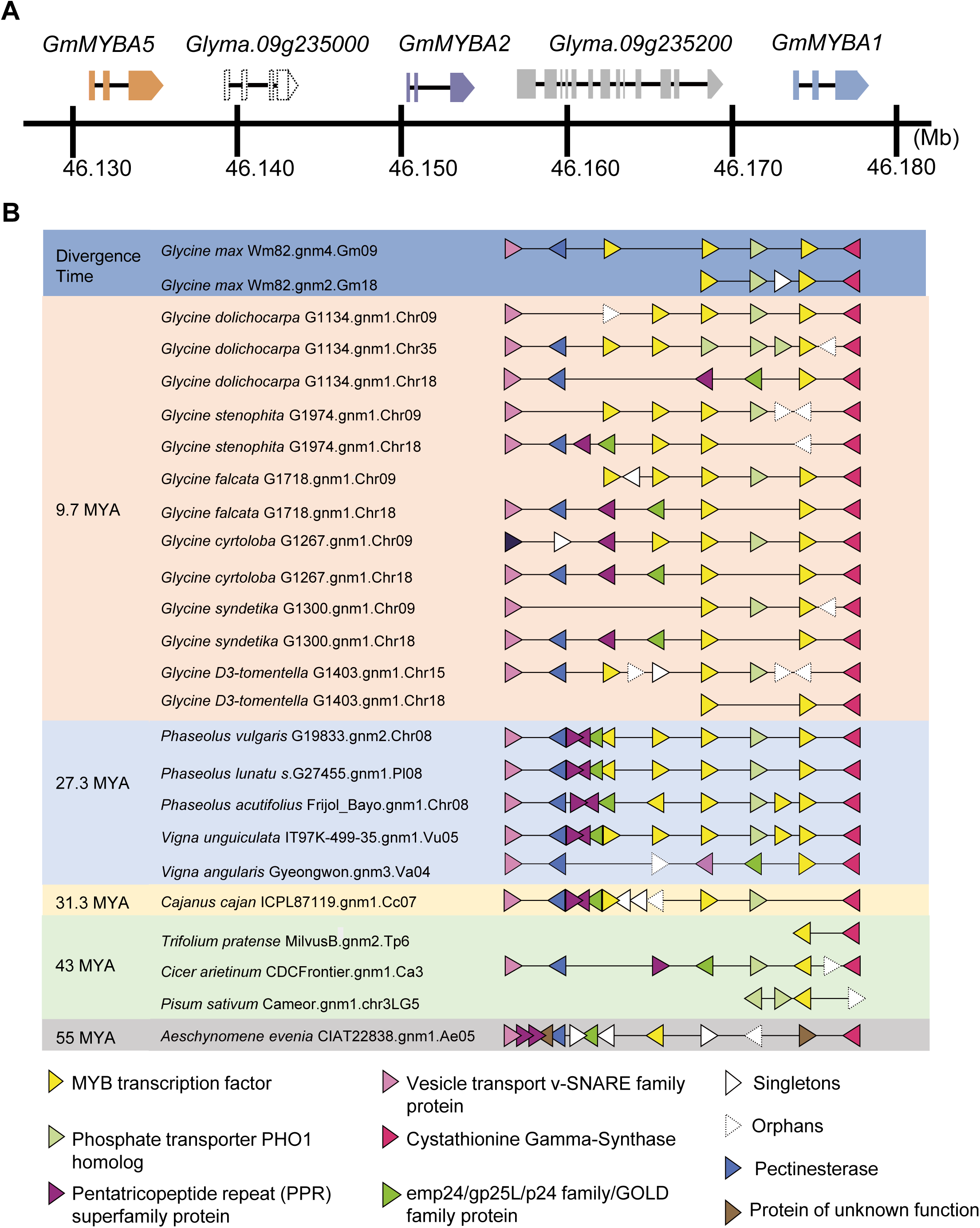
Micro-synteny analysis of the *MYB* gene cluster in legumes. (A) GmMYBA5 (Glyma.09g234900), GmMYBA2 (Glyma.09g235100), GmMYBA1 (Glyma.09g235300) and the pseudogene (Glyma.09g235000) are located in a cluster on chromosome 9. (B) Micro-synteny analysis of the MYB gene cluster in legumes. Each triangle represents a gene and the tip of the triangle indicates the direction of the gene. Genes belonging to the same family are depicted using the same color.

**Table 1.** Evolutionary distances of the three MYB genes within soybean and between soybean and common bean.

| MYB gene | Homologous gene in soybean or<br>common bean | Ka | Ks | $\omega$ (Ka/Ks) |
| --- | --- | --- | --- | --- |
| *Glyma.09G234900* (GmMYBA5) | *Phvul.008G038200* | 0.1823 | 0.3714 | 0.4907 |
| *Glyma.09G235100* (GmMYBA2) | *Phvul.008G038200* | 0.1776 | 0.3972 | 0.4471 |
| *Glyma.09G235300* (GmMYBA1) | *Phvul.008G038200* | 0.1843 | 0.4659 | 0.3956 |
| *Glyma.09G234900* (GmMYBA5) | *Glyma.09G235100* (GmMYBA2) | 0.0758 | 0.2030 | 0.3735 |
| *Glyma.09G234900* (GmMYBA5) | *Glyma.09G235300* (GmMYBA1) | 0.0940 | 0.2758 | 0.3407 |
| *Glyma.09G235100* (GmMYBA2) | *Glyma.09G235300* (GmMYBA1) | 0.0880 | 0.2721 | 0.3234 |
| *Phvul.008G038200* | *Phvul.008G038400* | 0.1740 | 0.4100 | 0.4244 |
| *Phvul.008G038200* | *Phvul.008G038500* | 0.2020 | 0.6456 | 0.3128 |
| *Phvul.008G038200* | *Phvul.008G038600* | 0.0356 | 0.0654 | 0.5444 |
| *Phvul.008G038400* | *Phvul.008G038500* | 0.0821 | 0.1414 | 0.5810 |
| *Phvul.008G038400* | *Phvul.008G038600* | 0.1904 | 0.5139 | 0.3705 |
| *Phvul.008G038500* | *Phvul.008G038600* | 0.2131 | 0.6243 | 0.3413 |

It is worth noting that our data revealed that the soybean specific whole-genome duplication (13 MYA) did not significantly increase the number of tandemly duplicated genes in this region. Instead, it led to an increase in the number of homoeologous genes within the duplicated region on chromosome 18 (Fig. 1B). Interestingly, while the homoeologous genes of *GmMYBA2* and *GmMYBA1* on chromosome 18 were retained, the homoeologous gene of *GmMYBA5* was lost, likely occurring subsequent to the whole-genome duplication event. Taken together, our data suggest that the *MYB* gene cluster originated through tandem duplication prior to the soybean specific whole-genome duplication and *GmMYBA5* has undergone greater functional divergence. Anthocyanins, as important metabolites, are often induced by environmental stresses in plants (Li et al., 2022b). The increased number of *MYB* genes in different legumes may represent an adaptive strategy for these plants to respond to diverse external environmental conditions.

### Tandemly duplicated *MYB* genes are putative anthocyanin synthesis regulators

The identification of *Glyma.09g235000* as a pseudogene (Gillman et al., 2011) prompted us to further investigate its function. To further confirm its role, we performed the ectopic expression of the coding sequence of *Glyma.09g235000* driven by the 35S promoter of cauliflower mosaic virus (CaMV) in *Arabidopsis thaliana*.

However, no phenotypic differences were observed in pigmentation between the transgenic plants and the wild types, indicating the loss of its regulatory function in anthocyanin synthesis (Supplementary Fig. S2). Therefore, we focused on the molecular function of the remaining three genes. Alignment of the amino acid sequences showed an 86.58% similarity among *GmMYBA5*, *GmMYBA2* and *GmMYBA1.* These three genes all contained the conserved R2 and R3 repeats, as well as the [D/E]Lx2[R/ K]x3Lx6Lx3R domain, which interacts with the R/B-like bHLH proteins (Supplementary Fig. S3). In addition, the conserved motif KPRPR[S/T] [F/L], which is important for anthocyanin activation, were present in all three genes (Stracke et al., 2001) (Fig. 2A). To elucidate the evolutionary relationship of these *MYB* genes, a phylogenetic tree was constructed using the protein sequences of known anthocyanin and proanthocyanin related MYB regulators from *Arabidopsis*, *Medicago* and *Vinifera.* The phylogenetic analysis classified these homologous genes into two major groups “anthocyanin activator” and “proanthocyanin activator”. Notably, all of the three *MYB* genes from the cluster fell within the “anthocyanin activator” group, indicating their potential roles in anthocyanin biosynthesis (Fig. 2B).

**Figure 2.**
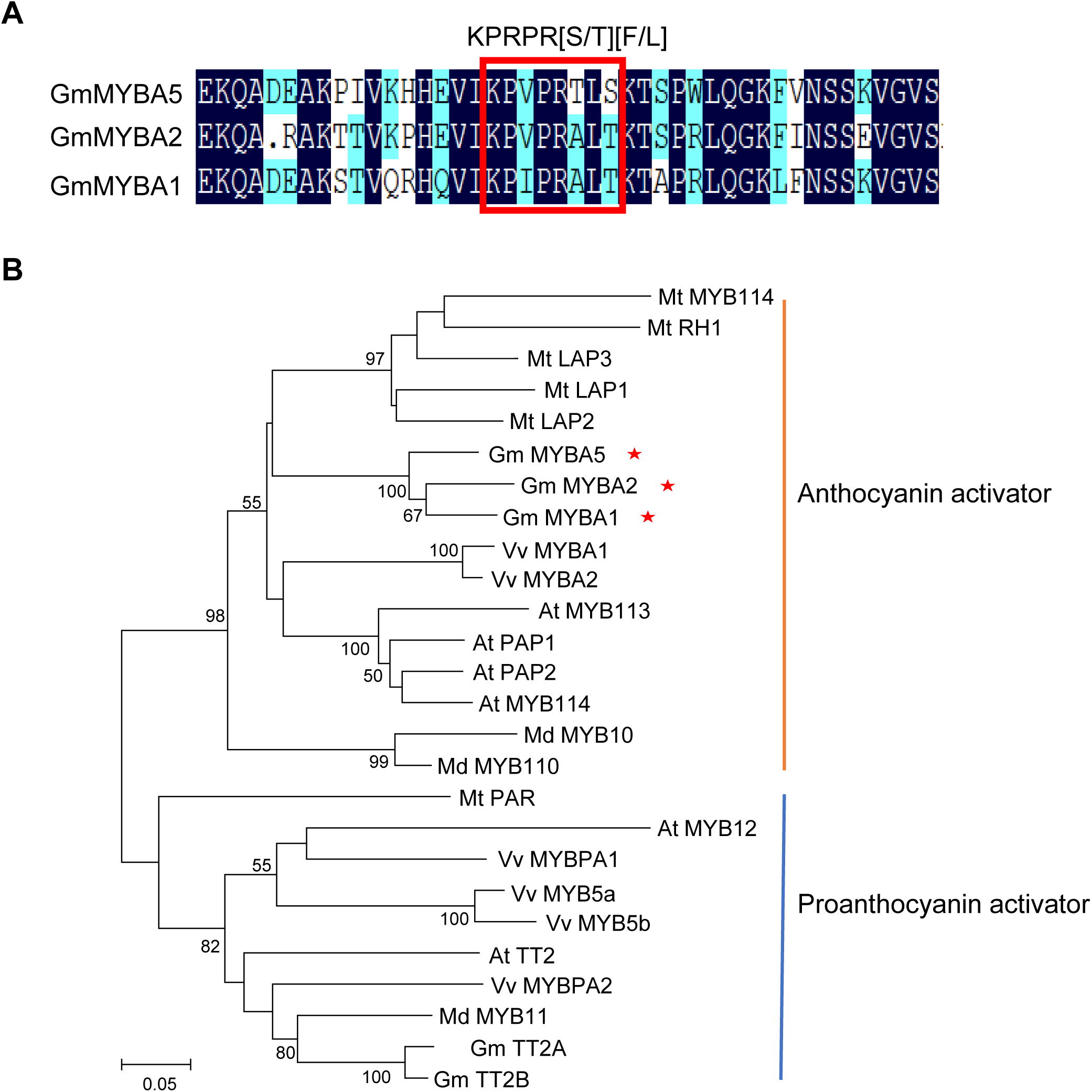
GmMYBA5, GmMYBA2 and GmMYBA1 are closely clustered and phylogenetically related in the soybean genome. (A) An amino acid sequence alignment of GmMYBA5, GmMYBA2 and GmMYBA1. The conserved KPRPR[S/T] [F/L] motif in subgroup 6 involved in anthocyanin regulation (activation) is highlighted in a red box. (B) Phylogenetic analysis of *MYB* genes involved in the anthocyanin and proanthocyanin pathways. The protein sequences were used to construct the phylogenetic tree using the neighbor-joining method in MEGA7, with 1000 bootstrap replicates. The numbers displayed at each node represent the bootstrap values that support the corresponding node, with vales above 50% from 1000 replicates being shown. The GenBank accession numbers corresponding to the MYB proteins are listed in Table S5.

### *GmMYBA5*, *GmMYBA2* and *GmMYBA1* are transcription factors that can activate anthocyanin biosynthesis

The subcellular distribution of GmMYBA5, GmMYBA2 and GmMYBA1 was investigated by expressing fusion protein of these genes with green fluorescent protein (GFP) in tobacco (*Nicotiana benthamiana*) leaf epidermal cells. The GFP fluorescent signals were observed specifically in the nuclei of all of the three genes (Fig. 3A), indicating that these genes are localized in the nucleus. To access their transcriptional activity, the full coding sequences of *GmMYBA5*, *GmMYBA2* and *GmMYBA1* were fused with the GAL4 DNA binding domain of yeast in the expression vector pGBKT7. The results showed that yeast transformants carrying these constructs were able to grow on medium lacking Trp, His and Ade, indicating that these three genes function as transcription factors with robust transcriptional activity (Fig. 3B). Furthermore, we introduced overexpression constructs containing the coding sequences of *GmMYBA5*, *GmMYBA2* and *GmMYBA1* driven by the 35S promoter of cauliflower mosaic virus (CaMV) into *Arabidopsis thaliana*. Compared to the wild type control, all five independent T_2_ transgenic lines exhibited purple pigment accumulation in various tissues, including seedlings, leaves, roots, and leaf veins (Fig. 3C and Supplementary Fig. S4). These findings demonstrate that *GmMYBA5*, *GmMYBA2* and *GmMYBA1* are nucleus-localized transcription factors with strong transcriptional activity that can activate the anthocyanin biosynthesis in vivo.

**Figure 3.**
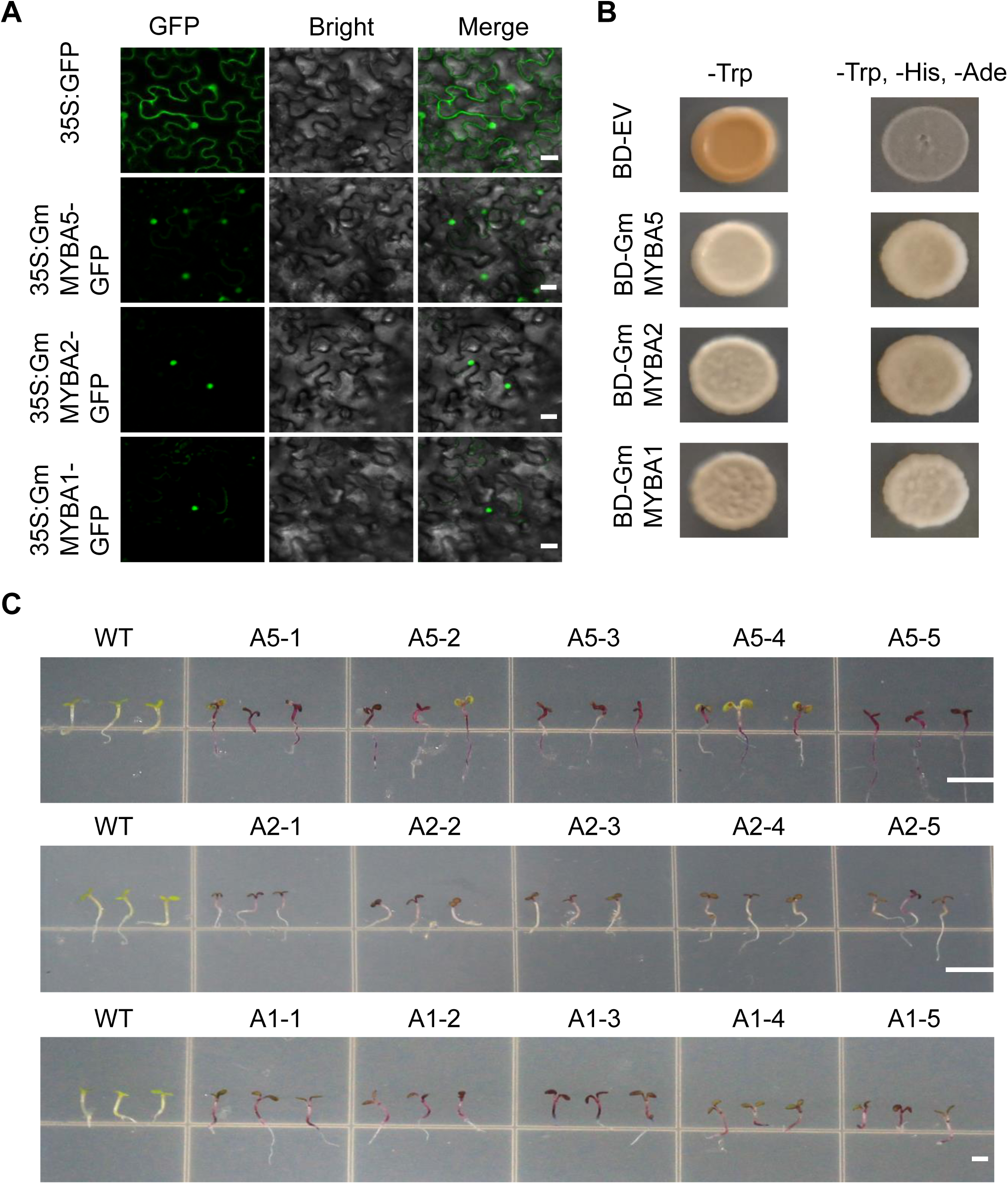
*GmMYBA5*, *GmMYBA2* and *GmMYBA1* function as transcriptional activators. (A) Subcellular localization of GmMYBA5, GmMYBA2 and GmMYBA1 in tobacco (*Nicotiana benthamiana*) leaf epidermal cells. Bars = 25 μm. (B) Transactivation activity assays in yeast demonstrated strong transactivation activity for GmMYBA5, GmMYBA2 and GmMYBA1. (C) Phenotype characteristics of *Arabidopsis* transgenic lines and wild type controls. WT, wild type; A5, 35S:*GmMYBA5*; A2, 35S:*GmMYBA2*; A1, 35S:*GmMYBA1*. Bars = 0.15 cm.

### Functional divergence of *GmMYBA5*, *GmMYBA2* and *GmMYBA1*

Previous studies have reported the specific expression pattern of *GmMYBA2* in the seed coat (Gillman et al., 2011). To validate this, quantitative reverse transcriptase (RT)-PCR analysis was performed on various soybean tissues, including stems, leaves, pods, and seeds. Consistent with previous findings, *GmMYBA2* was specifically expressed in seeds. Conversely, *GmMYBA5* was primarily expressed in vegetative tissues such as stems, leaves, and pods. In contrast, *GmMYBA1* displayed low expression levels restricted to leaves and stems (Fig. 4A). To further investigate the impact of ectopic expression of these three genes, transient overexpression of these three genes were conducted in *Nicotiana benthamiana* leaves. Leaves overexpressing *GmMYBA5*, *GmMYBA2* and *GmMYBA1* exhibited a noticeable anthocyanin-pigmented phenotype, and the anthocyanin content was significantly higher compared to the leaves with an empty vector control (Fig. 4B and C). Interestingly, the anthocyanin solution extracted from leaves overexpressing *GmMYBA5* displayed a brownish color instead of the purple color observed in leaves overexpressing *GmMYBA2* and *GmMYB1* (Fig. 4B). This observation suggests that the ectopic expression of *GmMYBA5*, *GmMYBA2* and *GmMYBA1* leads to the production of distinct metabolites in *Nicotiana benthamiana* leaves.

**Figure 4.**
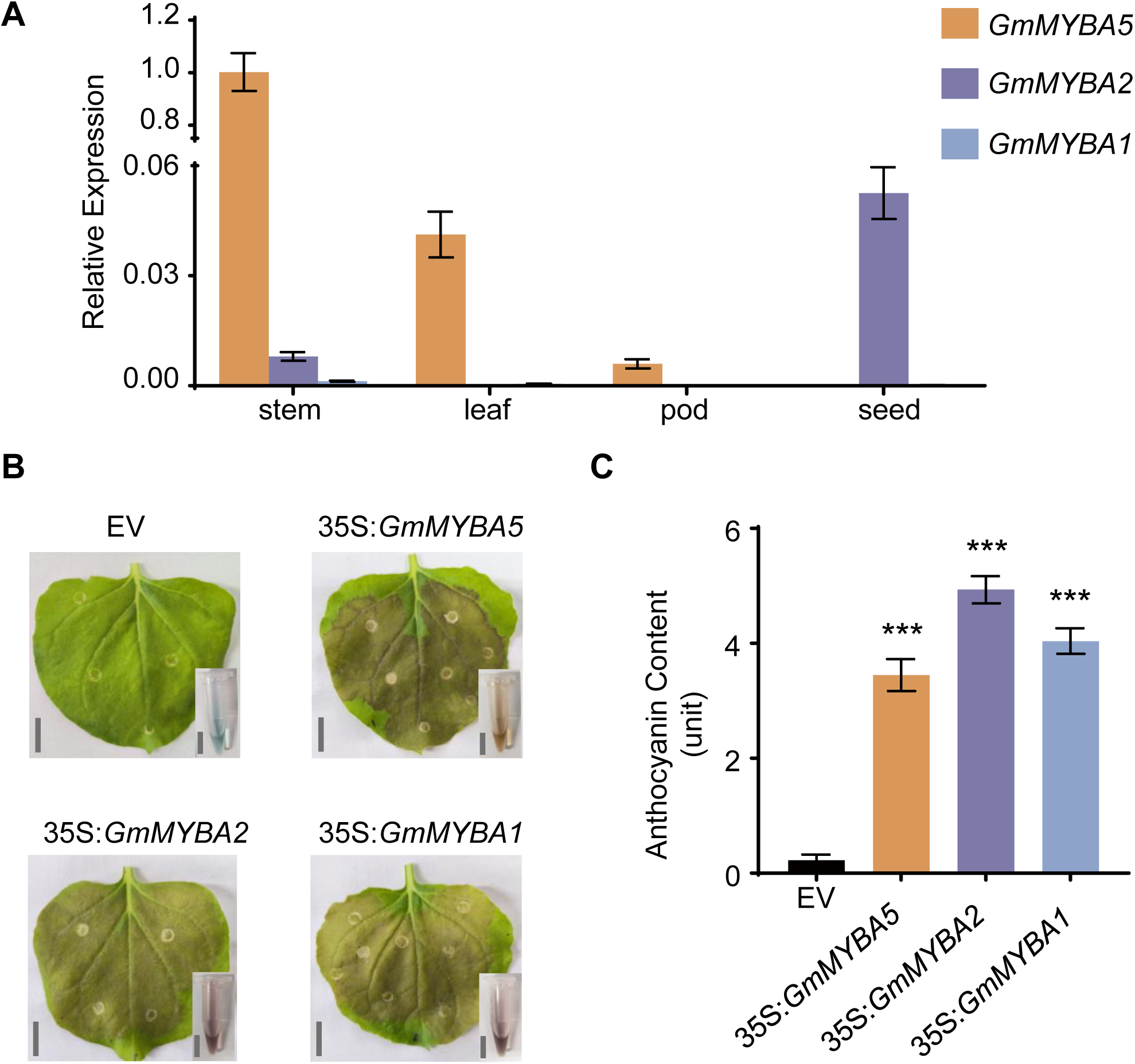
Functional divergence of the duplicated *MYB* genes. (A) Gene expression analysis of *GmMYBA5*, *GmMYBA2* and *GmMYBA1* in different tissues of soybean. (B) *Nicotiana benthamiana* leaves and total anthocyanin after the transient overexpression of *GmMYBA5*, *GmMYBA2*, *GmMYBA1* and the control with an empty vector (EV). Bar = 1 cm. (C) Overexpression of *GmMYBA5*, *GmMYBA2* and *GmMYBA1* induced significant enrichment of anthocyanins. The relative anthocyanin contents were quantified using the formula (A530-0.25×A657)/fresh weight, representing one anthocyanin unit. The data indicates the mean ± SD for three biological replicates. Statistical significance was determined using Student’s *t* test (***, p<0.001).

To elucidate the molecular basis underlying these different metabolites, domain swap experiments were performed. The DNA binding domains (BD; corresponding to R2R3 domain) and the activating domains (AD; corresponding to C-terminal domain) of *GmMYBA5*, *GmMYBA2* and *GmMYBA1* were exchanged, generating chimeric proteins (Fig. 5A). Our results showed that when the BD of *GmMYBA5* was fused with the AD of either *GmMYBA2* or *GmMYBA1*, the resulting chimeric proteins retained the phenotype of *GmMYBA5* (Fig. 5B). This indicates that the unique brownish metabolites produced by *GmMYBA5* is primarily influenced by its divergent R2R3 domains and potentially by its binding ability.

**Figure 5.**
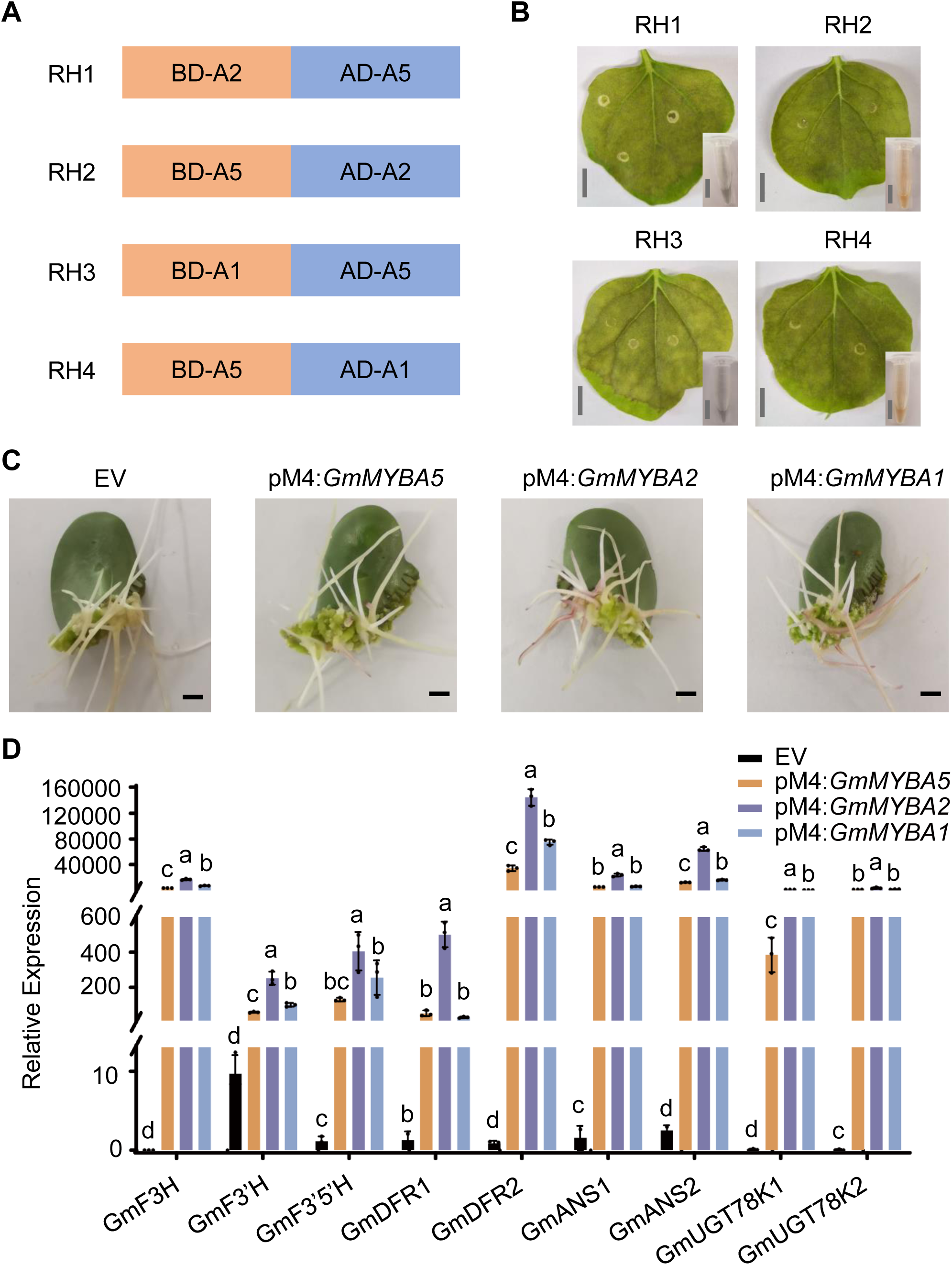
Molecular basis underlying the functional divergence of the *MYB* genes. (A) Chimeric proteins generated by fusing the DNA binding domains (BD) and activating domains (AD) from *GmMYBA5*, *GmMYBA2*, and *GmMYBA1*. (B) *Nicotiana benthamiana* leaves and total anthocyanin after the transient expression of different chimeric proteins. Bar = 1 cm. (C) Transgenic soybean hairy roots of Wm82 overexpressing empty vector (EV) and *GmMYBA5*, *GmMYBA2* and *GmMYBA1* with pM4. Bar = 0.25 cm. (D) Expression analyses of anthocyanin-related genes in transgenic soybean hairy roots of Wm82 overexpressing empty vector (EV) and *GmMYBA5*, *GmMYBA2* and *GmMYBA1* with pM4. Different lowercase letters indicate significant differences among groups based on Fisher’s Least Significant Difference (LSD) test at P < 0.05.

Additional experiments were performed in soybean hair roots to further investigate the function of the MYB genes. The highly expressed GmScreamM4 promoter (pM4) was utilized to replace 35S promoter of the plant expression vector pTF101, (Zhang et al., 2015), and the full coding sequences of *GmMYBA5*, *GmMYBA2* and *GmMYBA1* were inserted. These constructs were transformed into *Agrobacterium rhizogenes* K599, and soybean cotyledons were infected to generate transgenic soybean hairy roots. Transgenic soybean hairy roots carrying the three genes exhibited purple pigmentation in some roots, with the pM4:*GmMYBA5* transgenic soybean hairy roots displaying less pigmentation compared to the others (Fig. 5C). Expression analysis revealed that the expression levels of catalytic enzymes genes, including *GmF3H*, *GmF3’H*, *GmF3’5’H*, *GmDFR1*, *GmDFR2*, *GmANS1*, *GmANS2*, *GmUGT78K1*, and *GmUGT78K2*, were significantly upregulated in the overexpression lines, although to a lesser extent in *GmMYBA5* lines compared to the other lines (Fig. 5D). These results suggest that *GmMYBA5* has a lower capacity to activate catalytic genes involved in anthocyanin biosynthesis, resulting in reduced visible anthocyanin accumulation in roots. Overall, our data demonstrate that these tandemly duplicated *MYB* genes have undergone functional divergence, leading to tissue-specific expression patterns and differential activation abilities for anthocyanin biosynthesis.

### Metabolomics analysis of metabolites produced by *GmMYBA5*, *GmMYBA2* and *GmMYBA1*

To gain further insights into the functionality of the three *MYB* genes, we examined the downstream compounds in soybean hairy roots overexpressing pM4:*GmMYBA5,* pM4:*GmMYBA2* and pM4:*GmMYBA1*, as well as control roots with an empty vector, using Liquid Chromatograph Mass Spectrometer (LC-MS). In total, we identified 156 flavonoid compounds across all samples, with 80 (51.3%) belonging to flavones and flavonols, 30 (19.2%) classified as isoflavonoids, and 12 (7.7%) identified as anthocyanins (Fig. 6A and Supplementary Table S1). To ensure the reliability of the metabolite extraction and detection, we assessed the total ion flow diagram (TIC) of the mass spectrometry analysis and calculated the Pearson correlation coefficient for different quality control (QC) samples. Our data demonstrated a high correlation, indicating the repeatability of both metabolite extraction and detection (Supplementary Fig. S5A and B). Furthermore, we performed principal component analysis (PCA) and hierarchical clustering to evaluate the biological repeatability among the samples. The PCA score plot and hierarchical clustering heatmap clearly revealed that all samples were distinctly separated into the four expected groups, affirming the high quality and repeatability of the data (Supplementary Figs. S5C and S6).

**Figure 6.**
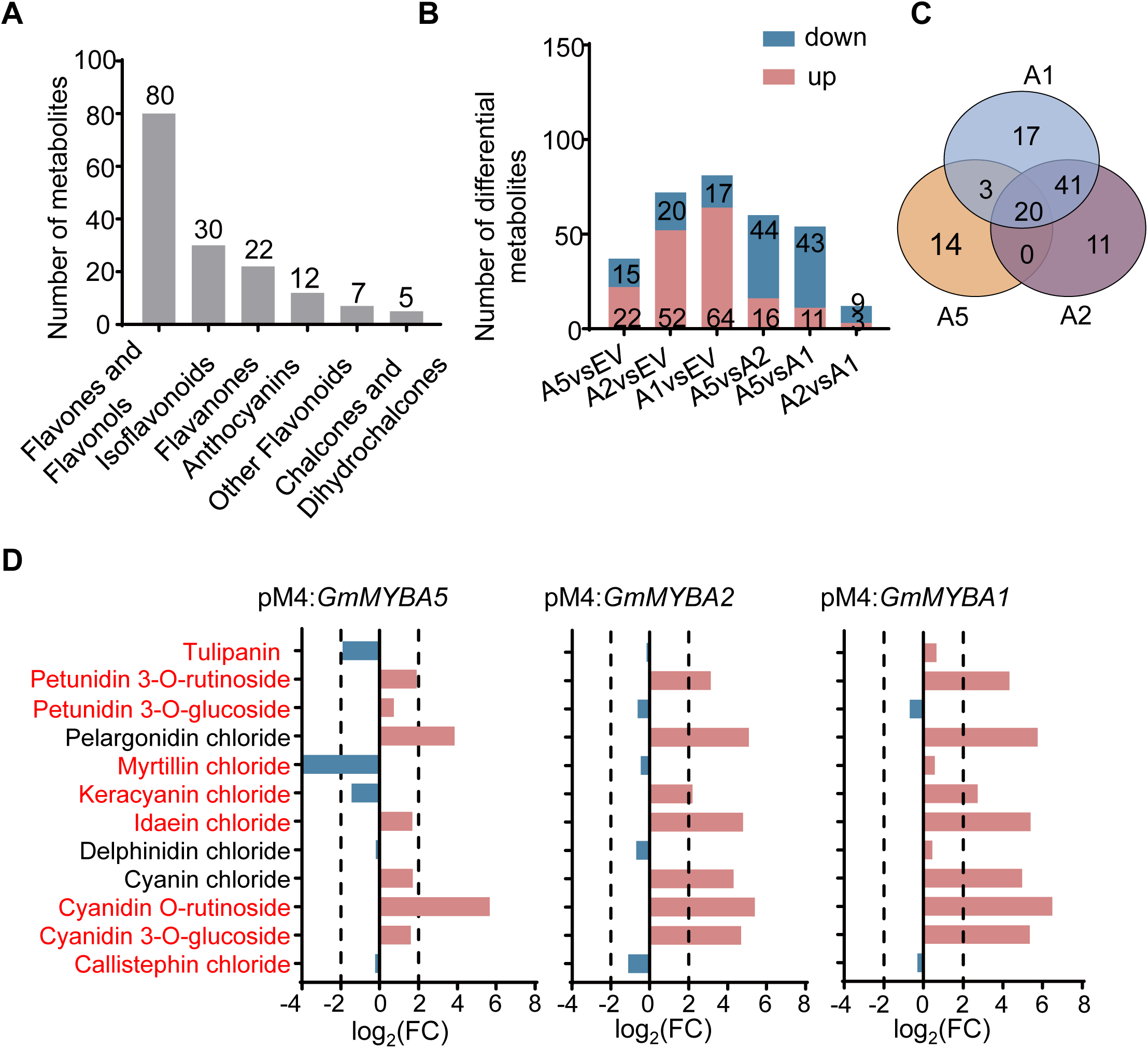
Metabolomics analysis of soybean hairy roots overexpressing the *MYB* genes revealed distinct types and abundances of downstream metabolites. (A) Classification and number of all flavonoids detected across the samples. (B) Number of differential metabolites between all comparisons. The numbers of up-regulated metabolites and down-regulated metabolites are shown in the histogram. EV, empty vector; A5, pM4:*GmMYBA5*; A2, pM4:*GmMYBA2*; A1, pM4:*GmMYBA1*. (C) Venn analysis depicting the overlapping and unique differential metabolites in hairy roots overexpressing *GmMYBA5, GmMYBA2* and *GmMYBA1.* A5, pM4:*GmMYBA5*; A2, pM4:*GmMYBA2*; A1, pM4:*GmMYBA1*. (D) Types and changes of anthocyanins detected in the overexpression lines of *GmMYBA5, GmMYBA2*, and *GmMYBA1.* Glycoside derivatives are highlighted in red.

The abundances of the 156 metabolic compounds were compared between the overexpression groups of the three genes and the control group with empty vectors, as well as within the overexpression groups. Differential metabolites were defined as those meeting the threshold criteria of p≤0.05 (Student’s *t* test) and |log_2_FC (Fold Change) |≥1. Our analysis revealed that the overexpression of *GmMYBA2* and *GmMYBA1* identified 72 and 81 differential metabolites, respectively, whereas *GmMYBA5* overexpression resulted in 37 differential metabolites (Fig. 6B). Among these differential metabolites, 22, 52, and 64 were upregulated, while 15, 20 and 17 were down-regulated when comparing the overexpression of *GmMYBA5*, *GmMYBA2* and *GmMYBA1*, respectively, with the empty vector (Fig. 6B, Supplementary Fig. S7A-F). Notably, the numbers of up-regulated flavonoids metabolites were greater than the number of down-regulated metabolites. Specially, out of the 37 differential metabolites driven by the overexpression of *GmMYBA5*, 22 (59.5%) were up-regulated, which is significantly lower than the numbers of the upregulated metabolites driven by the overexpression of *GmMYBA2* (72.2%) and *GmMYBA1* (79.0%) (Fig 6B). Among these up- and down-regulated metabolites, 20 (18.9%) were shared by all of the three genes (Fig. 6C and Supplementary Table S2). Furthermore, of the 72 differential metabolites driven by the overexpression of *GmMYBA2*, 61 (84.7%) were shared with *GmMYBA1*, suggesting potential redundancy in the regulation of flavonoid synthesis between *GmMYBA2* and *GmMYBA1* (Fig. 6B and C).

To further explore the regulatory role of these three *MYB* genes in anthocyanin biosynthesis, we focused on the analysis of 12 anthocyanins. These 12 anthocyanins encompassed cyanidin derivatives (idaein chloride, cyanin chloride, cyanidin O-rutinoside, cyanidin 3-O-glucoside, keracyanin chloride), delphinidin derivatives (myrtillin chloride, delphinidin chloride), pelargonidin derivatives (pelargonidin chloride, callistephin chloride), and petunidin derivatives (petunidin 3-O-rutinoside, petunidin 3-O-glucoside) (Fig. 6D). Our data showed that the majority of these 12 anthocyanins were upregulated in the overexpression lines of the three *MYB* genes. Specially, in the *GmMYBA1* overexpression line, 10 (83.3%) out of the 12 anthocyanins were upregulated (Fig. 6D). In contrast, *GmMYBA5* overexpression resulted in fewer and lower levels of anthocyanins in soybean hairy roots, which could explain the reduced pigmentation observed in these roots. This observation is consistent with the expression levels of catalytic enzymes genes (Fig. 6D). It is worth noting that glycoside derivatives constituted a significant proportion (40.91% and 44.19%) of the differential metabolites between *GmMYBA5* and *GmMYBA2*, as well as between *GmMYBA5* and *GmMYBA1* (Fig. 6D and Supplementary Table S3). These data suggest that *GmMYBA5* has partially lost its function in regulating the expression of genes involved in anthocyanin biosynthesis, resulting in reduced production of glycoside derivatives. On the other hand, *GmMYBA2* and *GmMYBA1* appear to have redundant functions in regulating the synthesis of flavonoids compounds including anthocyanins.

## Discussion

### MYB transcription factors are possible hotspots for tandem duplication

Tandem duplication represents one of the key processes by which the copy number of genes can be increased, leading to the emergence of new genetic resources during the course of evolutionary history in many organisms. Certain classes, such as *R* genes, are known to be prone to tandem duplication (Meyers et al., 2003; Innes et al., 2008).

It appears that *MYB* genes are also hotspots for tandem duplication in soybean, as well as in several other species (O’Neil et al., 2007; Wang et al., 2023). Previous studies have demonstrated that selection is relaxed on tandemly duplicated genes relative to non-tandemly duplicated genes. This relaxed selection may allow more transposon insertions or less efficient purging of transposons near these genes (Zhao et al., 2017). However, this does not seem to be the case for the three *MYB* genes examined in our study. Although these *MYB* genes have undergone functional divergence, they still retain the function of the ancestral gene in regulating anthocyanin biosynthesis. According to the gene dosage balance hypothesis, genes encoding proteins that interact with other proteins are more sensitive to changes in order to maintain the overall network functionality (Birchler and Veitia, 2021). Given that these *MYB* genes function as transcription factors, they likely interact with other proteins, which could explain their conserved function. Nevertheless, our study reveals a tissue-specific pattern for these gene (Fig. 4A), a characteristic commonly observed in many other tandemly duplicated genes (Zhao et al., 2017).

To understabnd the mechanism underlying MYB tandem duplication, we also examined the transposon sequences within the genic and flanking regions of the three *MYB* genes. Surprisingly, we found very few transposon sequences enriched in these regions. However, considering the high rate of transposon turnover in soybean, it is likely that the elements that contribute to the tandem duplication of the *MYB* genes have already been eliminated from the genome. Further investigation could involve examining other recently tandemly duplicated *MYB* gene clusters to assess the role of transposons in mediating the tandem duplication of *MYB* genes.

### Functional divergence of the *MYB* duplicated genes

Subfunctionalization proposes that duplicated genes originating from a common ancestor specialize in complementary functions to maintain the original function of their ancestral gene (Sandve et al., 2018). In our research, the tandemly duplicated *MYB* genes have undergone both pseudogenization and subfunctionalization. A putative pseudogene (*Glyma.09g235000*) structurally resembled the gene but lacked expression and function. Other genes within the *MYB* gene cluster acted as activators of anthocyanin synthesis but exhibited specialization in tissue-specific expression patterns, varied activation abilities, and diverse downstream metabolites (Fig. 7). Conserved MYB transcription factors contain activation domain in the C-terminal and A binding domain in the N-terminal, enabling them to regulate anthocyanins synthesis by forming MYB-bHLH-WD ternary complexes. (Wang et al., 2021). We have demonstrated that variations in the BD contribute to the divergent anthocyanin productions. The conserved bHLH-interacting motif [D/E]Lx2[R/ K]x3Lx6Lx3R is present in GmMYBA5, GmMYBA2 and GmMYBA1, albeit with two substitutions of L (Leucine) to M (Methionine) (Liu et al., 2015) (Supplementary Fig. S3). However, whether these three genes exhibit distinct interaction preferences for basic helix-loop-helix proteins, thereby resulting in the different metabolites, remains to be determined.

**Figure 7.**
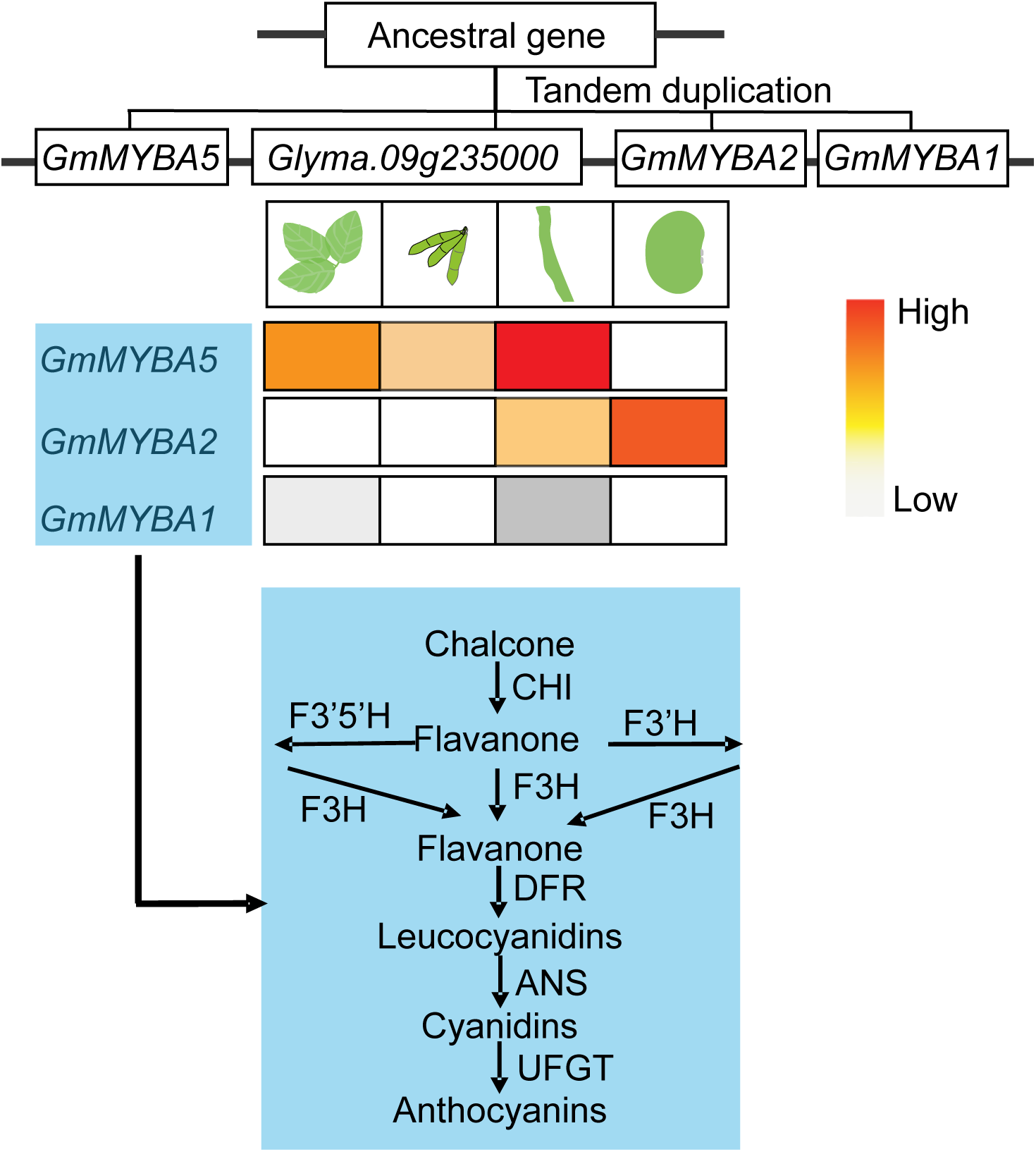
The model of the tandemly duplicated *MYB* genes functionally diverged in the regulation of anthocyanin biosynthesis pathway. The *MYB* genes following tandem duplication undergone functional divergence, leading to tissue-specific expression patterns and differential activation abilities for anthocyanin biosynthesis pathway. CHI, chalcone isomerase; DFR, dihydroflavonol-4-reductase; F3’H, flavonoid 3’-hydroxylase; F3’5’H, flavonoid 3’5’hydroxylase; F3H, flavanone 3-hydroxylase; DFR, Dihydroflavonol 4-reductase; ANS, anthocyanidin synthase; UFGT, UDP-flavonoid glucosyltransferase.

*GmMYBA2* plays a regulatory role in the late stage of anthocyanin biosynthesis by controlling enzymes such as flavonoid 3’5’hydroxylase, dihydroflavonol 4-reductase, anthocyanidin synthase and glucosyltransferase, which are involved (Gao et al., 2021). In contrast, overexpression of *GmMYBA5* led to reduced anthocyanin production, indicating a partial loss of its regulatory function in regulating catalytic enzymes genes. On the other hand, *GmMYBA2* and *GmMYBA1* exhibit functional redundancy in regulating anthocyanidin synthesis, although they display distinct expression patterns (Fig. 3A). Furthermore, the expression of *GmMYBA5* and *GmMYBA1* is generally induced by abiotic and biotic stresses, as inferred from publicly available RNA-seq libraries (http://ipf.sustech.edu.cn/pub/soybean/), suggesting their putative function in adapting to changing environment factors (Zhang et al., 2020) (Supplementary Fig. S8A-C). These results offer additional insights into the evolutionary trajectory and function divergence resulting from gene duplication in the regulation of plant secondary metabolites.

### Insights into the relationship of gene duplication and environmental adaptation

Gene duplication is an important force for genome evolution and environmental adaptation (Magadum et al., 2013; Chen et al., 2022b). Numerous studies have highlighted the occurrence of adaptive gene duplications in response to environmental stresses (Brown et al., 1998; Riehle et al., 2001; Hastings, 2007; James et al., 2008). In our investigation of the collinear regions of the MYB transcription factors cluster in legumes, we found that these orthologous *MYB* genes contribute to seed coat color in common bean and cowpea (García-Fernández et al., 2021; Herniter et al., 2018). The conservation of MYB transcription factors across species suggests their essential role in enabling efficient responses to environmental stresses (Saigo et al., 2020). The observed copy-number variation and functional divergence of *MYB* genes in legumes serve as a compelling example of adaptive gene duplications in the evolutionary history of legumes, which enhances our understanding of the role of gene duplications as a mechanism of adaptation.

### Anthocyanin can be used as a morphological marker

As Anthocyanin pigmentation can be identified by naked eyes, it has been employed as a visible marker in the maize haploid inducer system, facilitating the efficient selection of haploid embryos (Chen et al., 2022a). Selecting transgenic tissues and plants during plant genetic transformation is a laborious and time-consuming process. Recently, R2R3 type MYB transcription factors have been utilized as visible markers for the selection of transformed tissues and plants in various species (Zhang et al., 2019; Huang et al., 2021; Lim et al., 2022). Soybean hairy roots can be obtained by *Agrobacterium rhizogenes* mediated transformation, forming chimera plants with transformed hairy roots and untransformed shoots. In our study, we constructed plant expression vectors containing the GmScreamM4 promoter (pM4), which drove the expression of *GmMYBA2* and *GmMYBA1* in transgenic soybean hairy roots, leading to the manifestation of a purple color (Fig. 5C). The application of *GmMYBA2* and *GmMYBA1* as strong activators of anthocyanin biosynthesis in soybean holds promising prospects for their utilization as markers to facilitate the effective identification of transformed hairy roots.

## Materials and Methods

### Plant materials

The soybean cultivar Wm82 (Williams 82) was grown in the greenhouse at China Agriculture University. Plant tissues were collected at the reproductive growth stage R4 (full pod) and immediately frozen into liquid nitrogen before being stored at -80 °C. The *Arabidopsis thaliana* ecotype Columbia was used for genetic transformation. The transient ectopic expression material, tobacco (*Nicotiana benthamiana*), was grown in the incubator with the constant temperature of 25℃.

### Estimation of evolutionary distance and analysis of micro-synteny

To estimate evolutionary distance, we followed the method previously described (Zhao et al. 2017; Yin et al. 2022). In brief, the coding sequences of the homologous genes were aligned using ClustalW (Thompson et al., 1994) with default parameters, followed by manual inspection. Pairwise alignments of the homologous genes were performed to calculate Ka and Ks using the yn00 module under the PAML software (Yang, 2007). Micro-synteny analysis was conducted using the LegumeInfo database (https://www.legumeinfo.org/). The divergence time was obtained from the TimeTree database (https://timetree.org/).

### cDNA synthesis and quantitative real-time PCR analysis

Total RNA was isolated from soybean leaves, pods, stems and seeds by the StarSpin HiPure Plant RNA Mini Kit (GenStar). To eliminate any genomic DNA contamination, the StarScriptII First-strand cDNA Synthesis Mix With gDNA Remover (GenStar) was employed to synthesize the first-strand complementary DNA (cDNA), following the manufacturer’s instructions. Quantitative real-time PCR (qRT-PCR) analysis was performed using 2×RealStar Green Fast Mixture (with ROX II) (GenStar) according to the manufacturer’s protocol, and amplification was carried out using the ABI 7500 Real-time PCR system (Applied Biosystems, USA). The *Actin11* gene was used as the internal control. The qRT-PCR data were analysed using the 2^-ΔΔCt^ analysis method. Details of all the primers can be found in Supplementary Table S4.

### Analyses of phylogenetic relationship

Sequence alignment of *GmMYBA5*, *GmMYBA2* and *GmMYBA1* was performed by DNAMAN software version 8.0. A phylogenetic tree was constructed using the Neighbor-Joining method in the MEGA7 program. The statistical significance of individual nodes was assessed by bootstrap analysis with 1,000 replicates.

### Arabidopsis thaliana transformation

The full-length coding sequences of *GmMYBA5*, *GmMYBA2* and *GmMYBA1* cloned from Wm82 and the synthesized code sequence of *Glyma.09g235000* were introduced into the pTF101 vector. The coding sequences were driven by the 35S promoter of cauliflower mosaic virus (CaMV). The resulting constructs were then introduced into *Agrobacterium* strain GV3101 for transformation of *Arabidopsis thaliana* using the floral dip method (Bent, 2006). The presence of the constructs in the transgenic plants was confirmed by PCR, followed by the sequencing of the PCR fragment with specific primers. All primers used are listed in Supplementary Table S4.

### Subcellular localization

The full-length CDS of *GmMYBA5*, *GmMYBA2* and *GmMYBA1* were cloned from Wm82. These CDS sequences were fused in-frame with the GFP coding sequence and subsequently inserted into a vector through combinational joining. The resulting constructs were introduced into *Agrobacterium* strain GV3101 for infection of *Nicotiana benthamiana* leaves. The GFP fluorescent signals emitted by the tobacco leaf tissues were captured using a Zeiss confocal laser scanning microscope (Zeiss, Germany).

### Transcriptional activation activity assay

The transactivation activity assay was performed as previously described (Hou et al., 2022). In brief, the coding sequences of *GmMYBA5*, *GmMYBA2* and *GmMYBA1* were fused with the GAL4 DNA-binding domain (BD) in the plasmid pGBKT7. These constructs were subsequently transformed into the yeast strain AH109 according to the procedure previously described (Gietz et al., 2007). The yeast colonies were then patched onto SD/-Trp and SD/-Trp/-His/-Ade plates and incubated at 30°C for 3 days.

### Transient ectopic expression in *Nicotiana benthamiana*

The pTF-gene constructs were used to perform transient ectopic expression in *Nicotiana benthamiana*, follwoing the procedure described earlier in the subcellular localization section. After a 5-day incubation period, the phenotype of the infiltrated leaf materials was observed, and the extracted anthocyanins were measured.

### Soybean hairy root transformation

The 35S promoter of cauliflower mosaic virus (CaMV) in the plant expression vector pTF101 was substituted with the pM4 promoter of soybean. The plasmids were then transformed into *Agrobacterium rhizogenes* K599 for infection of soybean cotyledons to generate transgenic soybean hairy roots. The procedure for transformation and infection followed the method previously described with slight modifications (Kereszt et al., 2007; Guo et al., 2011).

### Anthocyanin extractions and measurements

Total anthocyanin contents were determined as previously described (Huang et al., 2018). Plant tissues were collected and subsequently ground into powders using liquid nitrogen. A total of 0.2 g powders was extracted in 1 ml of methanol containing 0.1% (v/v) HCl, followed by incubation on ice for 30 min. After centrifugation at 12,000 rpm for 10 min, the supernatant was measured at 530 nm (A530) and 657 nm (A657) using a spectrophotometer (SOPTOP, China). The relative content of anthocyanin was calculated using the formula (A530-0.25×A657)/fresh weight.

### Metabolomics analysis

Samples were prepared and metabolites were extracted following the protocols provided by Novogene Co., Ltd. (Beijing, China). The extraction solution was then injected into the LC-MS/MS system. LC-MS/MS analyses were performed using an ExionLC™ AD system (SCIEX) coupled with a QTRAP® 6500+ mass spectrometer (SCIEX) at Novogene Co., Ltd. (Beijing, China). The detection of the experimental samples using Multiple Reaction Monitoring (MRM) was based on the in-house database of Novogene. The generated data files from the HPLC-MS/MS were processed using the SCIEX OS Version 1.4 to integrate and correct the peaks. Sample normalization and significantly differential accumulated flavonoids were analyzed using MetaboAnalyst 5.0 (https://www.metaboanalyst.ca/). Samples were normalized by sum and the data were transformed using log10. The standard threshold criteria were set as follows: p≤0.05 (Student’s *t* test) and |log_2_FC |≥1. Principal component analysis (PCA) and hierarchical clustering were carried out using MetaboAnalyst 5.0 (https://www.metaboanalyst.ca/).

## Funding

This work was supported by the National Natural Science Foundation of China (grant nos. 32072089 and 31871708), the Recruitment Program of Global Experts and the Fundamental Research Funds for the Central Universities (2022TC021).

## Author Contributions

L.S. and M.Z. conceived and designed the project. R.M., Q.H., G.T. and J.L. performed the experiments and W.H., J.A., T.F., and J.H. analysed the data. R.M. wrote the manuscript. M.Z. and L.S. revised the paper. All authors read and approved of this manuscript.

## Conflicts of Interest

The authors declare that they have no conflicts of interest associated with this work.

